# Precise genome engineering in *Drosophila* using prime editing

**DOI:** 10.1101/2020.08.05.232348

**Authors:** Justin A. Bosch, Gabriel Birchak, Norbert Perrimon

**Author notes:** Corresponding authors: Justin A. Bosch and Norbert Perrimon, Harvard Medical School, 77 Avenue Louis Pasteur, Dept. of Genetics, NRB 336, Boston, MA 02115.

## Abstract

Precise genome editing is a valuable tool to study gene function in model organisms. Prime editing, a precise editing system developed in mammalian cells, does not require double strand breaks or donor DNA and has low off-target effects. Here, we applied prime editing for the model organism *Drosophila melanogaster* and developed conditions for optimal editing. By expressing prime editing components in cultured cells or somatic cells of transgenic flies, we precisely installed premature stop codons in three classical visible marker genes, *ebony*, *white*, and *forked*. Furthermore, by restricting editing to germ cells, we demonstrate efficient germ line transmission of a precise edit in *ebony* to ~50% of progeny. Our results suggest that prime editing is a useful system in *Drosophila* to study gene function, such as engineering precise point mutations, deletions, or epitope tags.

## Introduction

Genome editing is a versatile tool to study gene function in model organisms. For example, targeted gene deletions or point mutations can be used to disrupt gene function, create gain of function alleles, or model human disease mutations (Pickar-Oliver and Gersbach 2019). Furthermore, insertions can be used for gene tagging to detect or manipulate endogenous proteins (Vandemoortele *et al.* 2019). *Drosophila melanogaster* is an excellent model to study gene function because of its easy genetic manipulation, rich genomic resources, and conservation of cellular, developmental, and physiological processes with humans (Hales *et al.* 2015; Ugur *et al.* 2016). Importantly, genome editing tools involving clustered regularly interspaced short palindromic repeats (CRISPR) have been successfully applied in *Drosophila* to study gene function (Venken *et al.* 2016; Korona *et al.* 2017; Bier *et al.* 2018).

Prime editing is a recently developed CRISPR-based tool to engineer precise edits in the genome (Anzalone *et al.* 2019). Unlike precise editing using Cas9 and homology-directed repair (HDR), prime editing does not induce double strand breaks and does not require DNA template containing the edit. In addition, this method appears to have low off-target effects. Prime editing consists of two components, 1) a single guide RNA (sgRNA) with a 3’ extension encoding the edit, referred to as a prime editing guide RNA (pegRNA), and 2) a nickase mutant of Cas9 (nCas9^H840A^) fused with an engineered Moloney murine leukemia virus (M-MLV) reverse transcriptase (RT) enzyme, referred to as prime editor 2 (PE2). The pegRNA/PE2 complex induces a nick at the target site and reverse transcribes the edit from the pegRNA into the genome via the RT domain. Like Cas9/HDR, many types of precise edits are possible with prime editing, such as single base changes, deletions, or insertions.

While prime editing was originally developed in human cells (Anzalone *et al.* 2019), it has been quickly adopted in other organisms including mice (Anzalone *et al.* 2019; Liu *et al.* 2020; Surun *et al.* 2020) and plants (Butt *et al.* 2020; Chen 2020; Hua *et al.* 2020; Li *et al.* 2020; Lin *et al.* 2020; Tang *et al.* 2020; Veillet *et al.* 2020; Wang *et al.* 2020; Xu *et al.* 2020). Prime editing has been used to help correct disease mutations (Anzalone *et al.* 2019; Rousseau *et al.* 2020), introduce herbicide resistant alleles (Butt *et al.* 2020; Chen 2020; Hua *et al.* 2020; Xu *et al.* 2020), alter plant morphology (Butt *et al.* 2020), and model human disease mutations in organoids (Liu *et al.* 2020; Schene *et al.* 2020). Adapting and testing prime editing in additional organisms, particularly model systems, has great potential to improve the study of gene function. Here, we develop reagents and optimized conditions to conduct prime editing in *Drosophila*.

## Results

### Prime editing in cultured S2R+ cells

To initially test prime editing in *Drosophila*, we expressed prime editing components in cultured S2R+ cells by transfection. We used the S2R+ PT5 line (Neumuller *et al.* 2012) that constitutively expresses mCherry fluorescent protein and which has been previously used for CRISPR/Cas9 genome editing (Viswanatha *et al.* 2018). To express PE2 in S2R+ cells, we constructed two plasmids for constitutive expression. *pAct-PE2* expresses PE2 under the *Drosophila Actin5c* promoter (Supplemental Figure 1A), and *pUAS-PE2* (Figure 1A) expresses PE2 when used in combination with *pAct-Gal4* (abbreviated as *pAct>PE2*). This should result in high levels of PE2 expression due to signal amplification of the Gal4/UAS system (Brand and Perrimon 1993). In addition, to express pegRNAs in cells, we constructed an empty expression vector (*pCFD3-NS*) that lacks the sgRNA scaffold sequence (NS=No scaffold) (Figure 1B), which is a modified version of the sgRNA expression plasmid *pCFD3* (Port *et al.* 2014).

**Figure 1.**
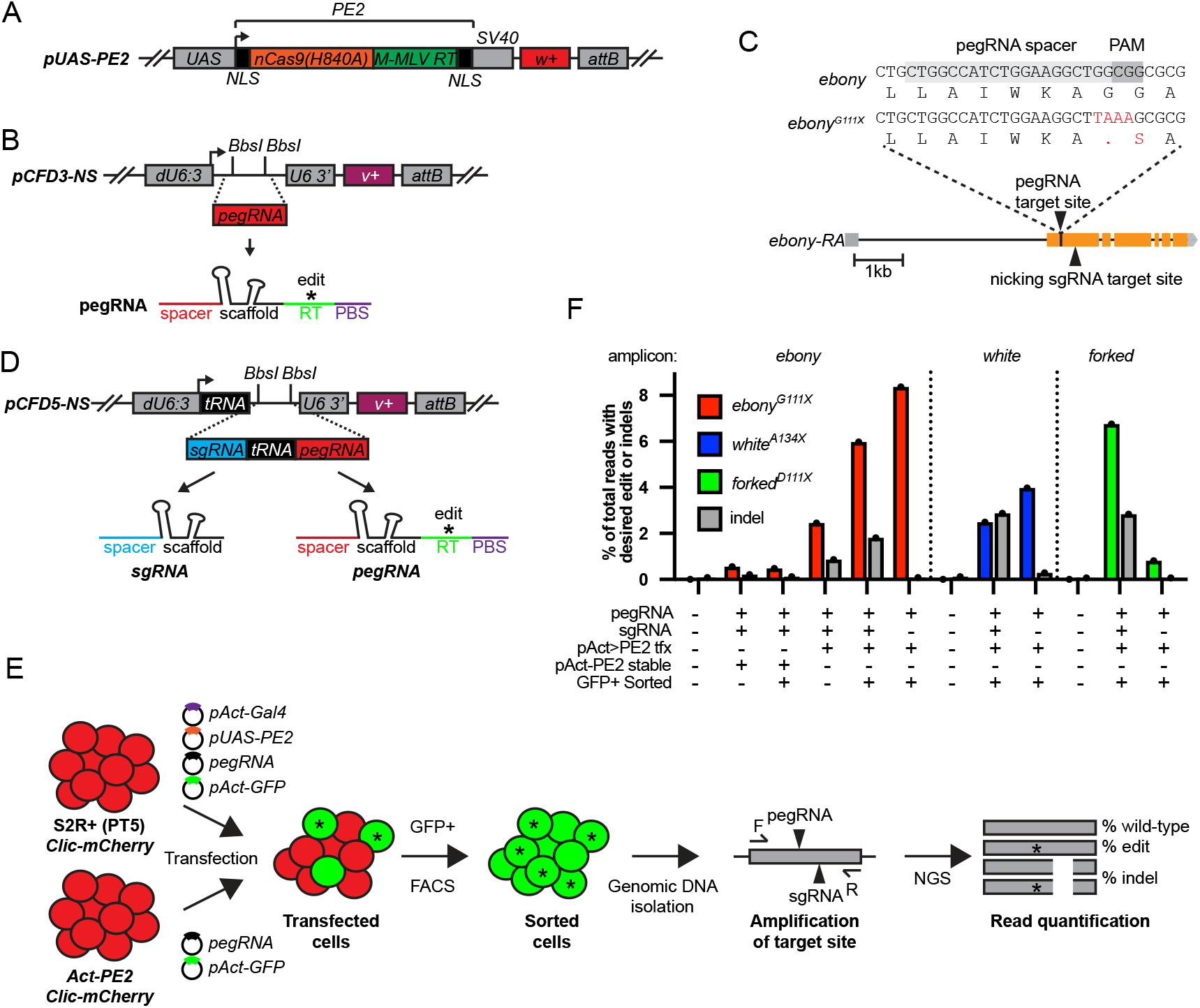
Prime editing in cultured S2R+ cells. **A.** Diagram of PE2 expression plasmid *pUAS-PE2*. *UAS*, *Upstream activating sequence*; NLS, Nuclear localization sequence; SV40, 3’ UTR; *w+*, *white+* rescue transgene; *attB*, phiC31 recombination site. **B.** Diagram of *pCFD3-NS* pegRNA expression plasmid. *BbsI* sites indicate cloning site for pegRNA encoding sequence. *dU6:3*, *U6* promoter; *U6* 3’, *U6* downstream region; *v+*, *vermillion+* rescue transgene. **C.** *ebony* genomic region showing target site and installed edit (*ebony*^*G111X*^). **D.** Dual sgRNA and pegRNA expression plasmid *pCFD5-NS*. tRNA, *D.m.* and *O.s.* Gly tRNA sequence. **E.** Schematic of S2R+ prime editing experiment. **F.** Quantification of precise editing and indels from S2R+ transfection experiments by amplicon sequencing. tfx, transfection.

First, we designed a pegRNA to insert a 23bp barcode (BC) sequence into the *ebony* gene (Supplemental File 1). This strategy was chosen to enable sensitive detection of insertion events by PCR. Four days after transfection of PE2 and pegRNA plasmids into PT5 cells, genomic DNA was collected and insertion-specific primers were used to amplify the putative insertion (Supplemental Figure 1B). Gel images of PCR products confirmed the presence of the *ebony*^*23bpBC*^ insertion using either *pAct-PE2* or *pAct>PE2* (Supplemental Figure 1C). To determine the insertion rate, we performed amplicon sequencing of the target region from transfected cells. Transfections using *pAct>PE2* resulted in an insertion efficiency of 0.42%, whereas transfections using *pAct-PE2* were substantially lower (0.006%) (Supplemental Figure 1D). Although our editing efficiencies were lower than reported in mammalian cells with an equivalent sized insertion (Anzalone *et al.* 2019), these initial results demonstrated that prime editing was possible in *Drosophila* S2R+ cells.

Next, we designed a pegRNA to introduce a premature stop codon in *ebony* (*ebony*^*G111X*^) (Figure 1C). In addition, we designed an sgRNA that nicks the non-edited DNA strand, since this approach, known as the Prime Editor 3 (PE3) system, can bias mismatch repair and boost editing efficiencies in mammalian cells (Modrich 2006; Chakraborty and Alani 2016; Anzalone *et al.* 2019). To simultaneously co-express a pegRNA and sgRNA, we constructed a dual expression vector called *pCFD5-NS* (Figure 1D). This vector uses tRNA processing to produce both pegRNA and sgRNA, and is a modified version of the multiplex sgRNA expression plasmid *pCFD5* (Port and Bullock 2016).

After transfecting PT5 cells with *pCFD5-PE3-ebony*^*G111X*^, *pAct>PE2*, and *pAct-GFP*, we isolated GFP+ cells using FACS and performed amplicon sequencing from their genomic DNA (Figure 1E). Under these conditions, precise editing efficiency of *ebony* was 6.0%. Furthermore, by comparing alternate conditions, we found that editing efficiency was ~2.5x lower without FACS enrichment and ~12x lower using a stable PE2 cell line (*Act-PE2*) (Figure 1E). Like in mammalian cells (Anzalone *et al.* 2019), the PE3 system caused a low percentage of insertions and deletions (indels) (0.86%) (Figure 1F). Finally, we compared editing efficiency using only a pegRNA (*pCFD3-PE-ebony*^*G111X*^). Unexpectedly, editing efficiency was slightly higher (8.4%) without a nicking sgRNA (Figure 1F). As expected, excluding the sgRNA reduced the frequency of indels to background levels.

To test prime editing at other genomic sites, we designed pegRNAs to introduce premature stop codons into *white* and *forked* (*white*^*A134X*^ and *forked*^*D111X*^), along with sgRNAs to nick on the non-edited strand (Supplemental Figure 1E). Editing efficiencies using both pegRNA and nicking sgRNA were roughly similar to *ebony*, producing 2.5% and 6.7% precise editing of *white* and *forked*, respectively. In addition, results with pegRNA only showed 4.0 and 0.8% precise editing of *white* and *forked*, respectively. Therefore, unlike *ebony* and *white*, *forked* editing efficiency was substantially improved by including a nicking sgRNA. In conclusion, using optimized prime editing conditions, we demonstrate precise editing efficiencies in S2R+ cells of ~4-8%.

### Prime editing in vivo

To test prime editing in vivo, we performed crosses between PE2 and pegRNA expressing transgenic flies. This strategy has been used with Cas9 (Bier *et al.* 2018), and Cas12a (Port *et al.* 2020a) to edit somatic and germ cells, and it is generally associated with higher editing efficiencies than embryo injection. To express PE2 in vivo, we generated *UAS-PE2* transgenic flies, which express PE2 when crossed with a Gal4 driver line (Figure 2A). In addition, we generated transgenic flies expressing pegRNAs to introduce premature stop codons into *ebony*, *white*, and *forked*. These genes/edits were chosen to enable easy identification of mutant flies with body phenotypes. In addition, transgenic pegRNA flies were created using the same plasmids validated in S2R+ cells (*pCFD3-PE-gene*^*edit*^ and *pCFD5-PE3-gene*^*edit*^).

**Figure 2.**
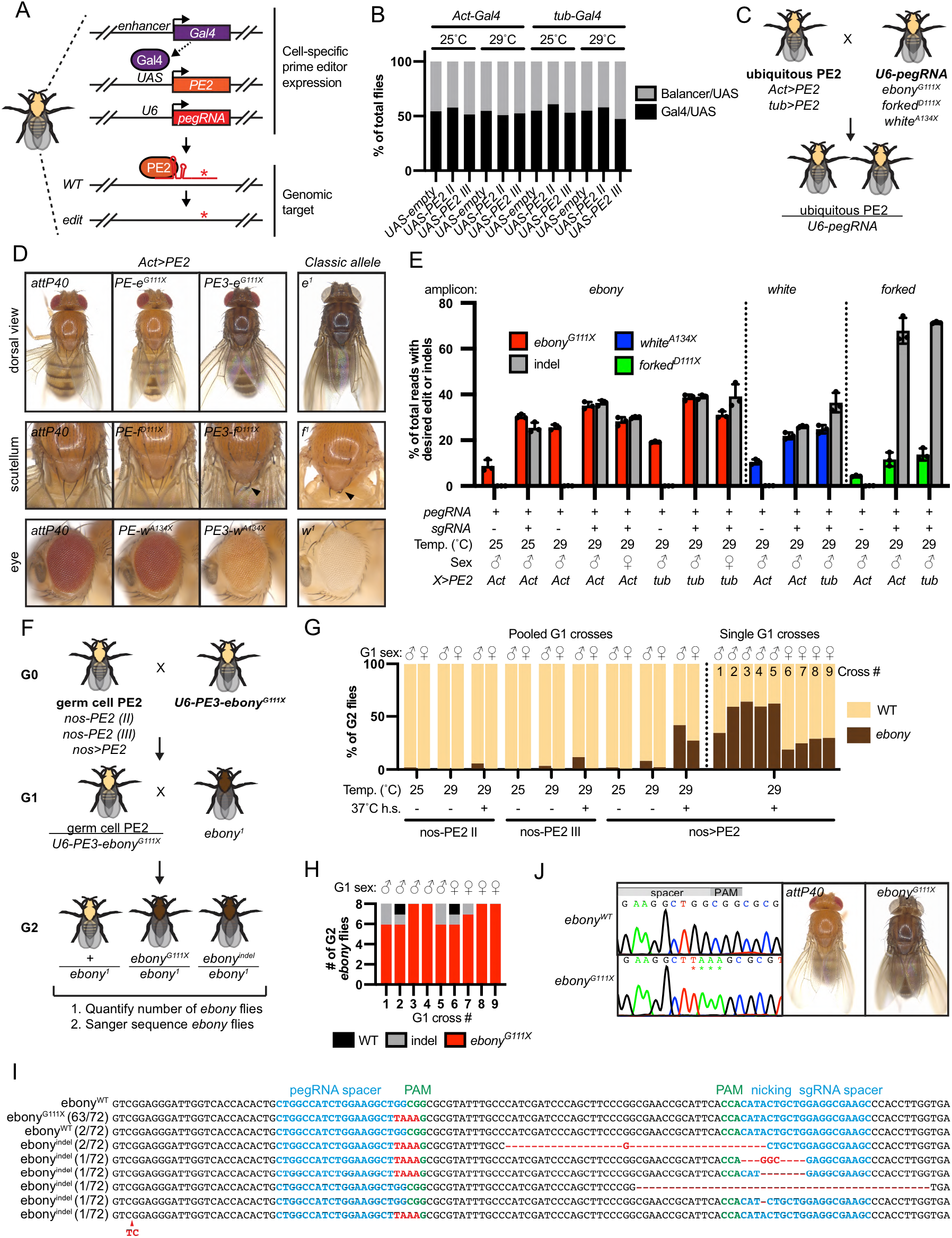
Prime editing in flies. **A.** Schematic of transgenic expression of prime editing components in flies and editing at an endogenous locus. Enhancer-specific Gal4 directs the spatial and developmental timing of PE2 expression. **B.** Quantification of adult fly viability from ubiquitous PE2 expression during developmental stages and raised at either 25°C or 29°C. *Act-Gal4/CyO* or *tub-Gal4/TM3* were crossed with *UAS-PE2* (Chr. II), *UAS-PE2* (Chr. III), or *UAS-empty* (negative control), and the percentage of progeny with or without the balancer was calculated. Number of flies scored from left to right = 748, 687, 655, 157, 267, 202, 294, 413, 226, 131, 277, 238. **C.** Schematic of genetic crosses between ubiquitous PE2 and pegRNA transgenic flies. **D.** Images of adult flies with somatic editing using *Act>PE2*. Views of the dorsal side of whole adults (top), scutellum (middle), and eye (bottom). Negative control is *attP40* and classical loss of function allele shown on right. Females shown for editing of *ebony* and *forked*, males shown for *white* editing. *e*^*1*^ = *w*^*1*^;; *TM3,e*^*1*^/*TM6b,e*^*1*^, *f*^*1*^ = *y*^*1*^, *w*^*1*^, *f*^*1*^. **E.** Quantification of precise somatic editing and indel percentage in adult flies by amplicon sequencing. Error bars show mean with SD. N=3 adult flies. **F.** Schematic of two generation genetic crosses between germ cell PE2 and pegRNA flies. **G.** Quantification of adult cuticle pigmentation (WT vs. *ebony*) in G2 flies for three temperature conditions. Sex of G1 parent(s) is indicated above graph. For the pooled crosses (10 G1 flies each, left of dotted line), the number of G2 flies analyzed was (left to right) 453, 518, 574, 413, 702, 405, 514, 454, 514, 405, 376, 493, 557, 492, 510, 562, 471, 481. For single fly G1 crosses (right of dotted line), the number of G2 flies analyzed was (left to right) 209, 109, 76, 139, 176, 104, 147, 275, 222. **H.** Quantification of Sanger sequencing analysis of individual G2 flies from single G1 crosses. Eight G2 progeny were analyzed for each of the nine G1 crosses. Sex of G1 parent is indicated above graph. **I.** Sequence of the *ebony* target site in 72 mutant *ebony* G2 flies. **J.** Sequence chromatogram (left) and image (right) of wild-type and *ebony*^*G111X*^ homozygous adult flies.

Many groups have reported toxicity in *Drosophila* from expression of Cas9 (Huynh *et al.* 2018; Poe *et al.* 2019; Port *et al.* 2020b) and Cas13 (Buchman *et al.* 2020). To test for toxicity from PE2 expression, we crossed *UAS-PE2* to two ubiquitous Gal4 drivers (*Act-Gal4* and *tub-Gal4*) and analyzed the resulting progeny (abbreviated as *Act>PE2* and *tub>PE2*). *Act>PE2* and *tub>PE2* larvae, pupae, and adults were morphologically normal (not shown). Furthermore, the observed number of *Act>PE2* and *tub>PE2* adult progeny was similar to negative control crosses when raised at 25°C or 29°C, and when using two different *UAS-PE2* transgenes (Figure 2B). Finally, *Act>PE2* and *tub>PE2* flies were fertile and could be propagated as a stock. Therefore, ubiquitous expression of PE2 does not result in obvious toxicity in flies.

Next, we crossed *Act>PE2* or *tub>PE2* to transgenic pegRNA lines and analyzed progeny for evidence of editing in somatic cells (Figure 2C). Crosses involving expression of a single pegRNA (*pCFD3-PE-gene*^*edit*^) resulted in progeny that were wild-type in appearance (Figure 2D, Supplemental Figure 2A). In contrast, somatic editing using the PE3 system (*pCFD5-PE3-gene*^*edit*^) resulted in progeny with mutant phenotypes similar to classical alleles (Figure 2D, Supplemental Figure 2A). In all cases, mutant phenotypes appeared slightly more severe at 29°C compared to 25°C (not shown). To determine the type and frequency of DNA changes at target sites, we performed amplicon sequencing from single adult fly genomic DNA. For *ebony*, *forked*, and *white,* precise editing efficiency using *Act>PE2* was highest with the PE3 system, resulting in 35.2%, 11.6%, and 21.9% reads, respectively, with the intended edit (Figure 2E). Comparable results were obtained using *tub>PE2* (Figure 2E). In addition, editing of *ebony* using *Act>PE2* was higher at 29°C than 25°C, but slightly lower in females compared to males (Figure 2E). The PE3 system led to a significant percentage of indels at the target site, with an exceptionally high percentage for *forked* (67.9%). Since both the precise edit and frameshift indels would cause loss of gene function, our sequencing results explain the strong mutant phenotypes when using the PE3 system in somatic cells.

Adapting prime editing to the germ line could enable the creation and propagation of edited fly stocks. To accomplish this, we generated transgenic flies with PE2 under the control of the germ cell-specific *nanos* (*nos*) promoter, either as a single transgene (*nos-PE2*) (Supplemental Figure 2B), or by combination of *nos-Gal4* with *UAS-PE2* (*nos>PE2*) (Figure 2A). We crossed *nos-PE2* or *nos>PE2* to *pCFD5-PE3-ebony*^*G111X*^ to generate G1 progeny with editing components expressed in germ cells (Figure 2F). Next, pools of 10 G1 progeny were crossed with *ebony*^*1*^ and the percentage of mutant *ebony* G2 progeny (*ebony*^*mut*^/*ebony*^*1*^) was calculated. Using this assay, we compared *nos-PE2* (two separate insertions) vs. *nos>PE2*, three temperature conditions (25°C, 29°C, and 29°C with 37°C heat shocks (hs)), and male vs. female germ line editing. We observed the highest transmission rate (42.2%) of *ebony* mutations from the G1 male germ line using *nos>PE2* and raising G1 animals at 29°C+hs (Figure 2G). Furthermore, single fly G1 crosses using *nos>PE2* and 29°C+hs produced similar results to pooled G1 crosses (Figure 2G) and 9/9 (100%) G1 flies were founders for mutation of *ebony*.

Next, we sequenced the *ebony* target site from 72 mutant *ebony* G2 progeny (Figure 2G). For each single fly G1 cross, the number of mutant *ebony* G2 flies with a correct edit ranged from 75% to 100% and was similar for male and female G1 crosses (Figure 2H). Combining sequencing results from all single fly G1 crosses and both G1 sexes, 63/72 (88.0%) of mutant *ebony* G2 flies had the desired edit, 7/72 (9.7%) had a frameshift indel, and 2/72 (2.8%) had wild-type sequence (Figure 2I). Taking the average frequency of mutant *ebony* G2 flies from single fly G1 crosses (56.2% from G1 males, 26.0% from G1 females) and multiplying by the frequency of *ebony* mutant flies with the *G111X* edit (88.0%), we estimate that male and female founders on average transmit the desired edit to 49.5% and 22.9% of progeny, respectively. Finally, homozygous *ebony*^*G111X*^ flies exhibited dark body pigment (Figure 2J) and could be propagated as a viable stock (not shown). These results demonstrate that prime editing is effective for engineering precise genomic edits in the *Drosophila* germ line.

## Discussion

Currently, precise genome editing in *Drosophila* is performed by CRISPR/Cas9 and homology directed repair (HDR) (Bier *et al.* 2018). HDR enables a wide variety of edits, yet is a relatively low-efficiency process, and a number of unintended side-effects have been documented, such as off-target mutations (Carroll 2013), imprecise integration of the donor DNA (Skryabin *et al.* 2020), or genome rearrangement (Ledford 2020). In addition, HDR is not as useful for tissue-specific editing because HDR events only occur in dividing cells. Furthermore, molecular cloning of donor constructs can be technically challenging and time-consuming.

Prime editing has the potential to address some of these limitations. PE2 uses a nickase mutant of Cas9 (H840A) that induces single strand breaks, which are known to decrease undesired genome changes and increase HDR:indel ratios (Maizels and Davis 2018; Anzalone *et al.* 2019). In addition, prime editing does not require cell division and functions in post-mitotic cultured cells (Anzalone *et al.* 2019). pegRNAs contain both targeting sequence and edit template and are simple to generate, thus facilitating multiple editing experiments in parallel. Furthermore, transgenic pegRNAs enable temporal and spatial control of precise editing, similar to transgenic sgRNAs used for CRISPR/Cas9 knockout (Kondo and Ueda 2013; Port *et al.* 2014; Meltzer *et al.* 2019; Poe *et al.* 2019; Port *et al.* 2020a). Generating transgenic pegRNA fly lines takes ~1 month, and thus delays germ line editing experiments compared to direct injection of genome-editing components into embryos. Injecting pegRNA plasmids or synthesized pegRNAs into PE2-expressing embryos, similarly to what is commonly done for Cas9-based HDR, could speed up the recovery of edited strains, but this approach remains to be tested for prime editing. One important caveat is that prime editing is currently limited to small (<100bp) edits that are identified by molecular assays (e.g. PCR).

Precise editing efficiencies in S2R+ cells were ~4x lower than in mammalian cells, and nicking sgRNAs (PE3 system) did not always increase efficiency. It is not clear if this is due to biological differences (e.g. DNA repair pathways) or technical differences (e.g. transfection method, promoter use, temperature) between these two culture systems. Further optimization of prime editing will likely improve its efficiency in cultured *Drosophila* cells. Regardless, our results suggest that prime editing can be used as a tool to generate edited S2R+ cells lines. Furthermore, pegRNAs could be stably integrated in S2R+ cells and used for pooled screening, as has been done with Cas9/sgRNAs (Viswanatha *et al.* 2018).

Ubiquitous PE2 and pegRNA expression in whole animals led to editing efficiencies of 10-40% for *ebony*, *white*, and *forked*. Although nicking sgRNAs led to higher editing frequencies, they also caused frequent indels (26-68%), which presumably contributed to the robust loss of function phenotypes we observed. Conversely, single pegRNAs did not cause obvious mutant phenotypes despite evidence of precise editing (4-26%). Therefore, unlike existing transgenic crossing techniques for somatic knockout (Port *et al.* 2014; Port and Bullock 2016; Meltzer *et al.* 2019; Poe *et al.* 2019; Port *et al.* 2020a; Port *et al.* 2020b), we were unable to install a precise edit in the majority of cells in the fly using ubiquitous expression of prime editing components. Nevertheless, some applications may be compatible with our reported somatic editing efficiencies, such as screening edits that drive tumorigenesis or affect cell competition.

By restricting expression of PE2 to germ cells, we demonstrated efficient transmission of a precise edit (*ebony*^*G111X*^) from transgenic founder flies to progeny. 100% of founder flies transmitted the *ebony*^*G111X*^ edit, with 49.5% of progeny from male founders inheriting the allele. This transmission rate is comparable to, if not higher than, using HDR and embryo injection to install similarly sized edits (Gratz *et al.* 2014; Port *et al.* 2014; Ge *et al.* 2016; Levi *et al.* 2020) and facilitates molecular screening of a small number of progeny. Similar to S2R+ and somatic cells, transmission rate was increased using Gal4/UAS-based PE2 expression and higher temperature, respectively. Further manipulating this temperature sensitivity will be useful to optimize germ cell editing. It will also be important to determine the generality of this method by test editing of additional genes, especially essential genes.

Currently, designing an effective pegRNA for precise editing is less straightforward than for sgRNAs. We deliberately selected pegRNA spacer sequences based on previously validated sgRNAs (see methods), but this might have led to better than average editing efficiency. The recent introduction of online tools have made pegRNA design easier, with options to optimize GC content and RNA stability (Chow *et al.* 2020; Hsu *et al.* 2020). When possible, we recommend testing editing efficiency in S2R+ cells before proceeding in vivo. While amplicon sequencing produces high quality quantitative data, there are faster and cheaper molecular assays such as the Dinucleotide signaTurE CapTure (DTECT) (Billon *et al.* 2020) or Tracking of Indels by Decomposition (TIDE) (Sentmanat *et al.* 2018).

In summary, we have developed genetic tools to express prime editing components in *Drosophila*, and optimized conditions for efficient editing in cultured cells and in vivo. By designing/cloning a pegRNA and optional sgRNA, *Drosophila* researchers can generate a wide variety of precise genome modifications such as point mutations, epitope tag insertions, or deletions. Furthermore, the ability to use prime editing in the fly germ line makes it useful to create custom fly strains for gene function analysis. Since CRISPR-based tools are continually engineered for optimal efficiency or new functions, it is likely that future variant prime editor systems will improve this method in *Drosophila*. Finally, the tools and optimized conditions we developed for prime editing in *Drosophila* may be useful in other model organisms.

## Acknowledgements

We thank Rich Binari for general lab assistance and help with fly genetics (particularly during the COVID-19 shutdown), TRiP and DRSC for help generating transgenic flies, Ram Viswanatha for sharing unpublished reagents and general discussions, Gillian Millburn for discussions on pegRNA transgene nomenclature, Cathryn King for general lab assistance, Cooper Cavers for help isolating transgenic flies, Jorden Rabasco for help with molecular cloning, and Ben Ewen-Campen, Jonathan Zirin, and Thai LaGraff for comments on the manuscript. J.A.B. was supported by the Damon Runyon Foundation and the “Training Grant in Genetics” T32 Ruth Kirschstein-NRSA institutional research training grant funded through the NIH/NIGMS. This work was also supported by NIH grants R24OD01984, R24OD030002 and P41GM132087. N.P. is an investigator of the Howard Hughes Medical Institute.

## Author contributions

Conceptualization and methodology, J.A.B.; Formal analysis J.A.B. and G.B.; Investigation, J.A.B. and G.B.; Writing – Original Draft, J.A.B.; Writing – Review & Editing, J.A.B., G.B., N.P.; Visualization, J.A.B.; Supervision, J.A.B., N.P.; Funding acquisition, J.A.B., N.P.

## Methods

### pegRNA and sgRNA design

pegRNA spacer sequences were selected based on previously validated sgRNA target sites for *ebony* (Port *et al.* 2015), *white* (Kondo and ueda 2013), and *forked* (Port and Bullock 2016). 13bp was used for the pegRNA prime binding site (PBS). For the reverse transcribed (RT) region, we used either a 34bp (*ebony*^*23bpBC*^) or 18bp (*ebony*^*G111X*^, *white*^*A134X*^, *forked*^*D111X*^) region. In all of our pegRNA designs, the pegRNA PAM is disrupted by the edit. Nicking sgRNAs were designed to nick the DNA strand opposite to the pegRNA-nicked strand within 40-90 bp of the pegRNA nick (*ebony*^*G111X*^: +57, *white*^*A134X*^: +70, *forked*^*D111X*^: +57). See Supplemental File 1 for pegRNA and sgRNA sequences. See Supplemental File 3 for additional pegRNA and sgRNA design parameters.

### Plasmid cloning

Plasmid DNAs were constructed and propagated using standard protocols as follows. PCR fragments were amplified using Phusion polymerase (New England Biolabs M0530). Plasmids were digested with restriction enzymes at 37°C for 2-16hrs. Linearized plasmid and PCR fragments were gel purified using QIAquick columns (28115, Qiagen). Inserts and backbones were assembled using Gibson assembly (New England Biolabs E2611) or T4 ligation (New England Biolabs M0202). Gateway-compatible expression and entry vectors were recombined using LR Clonase II (ThermoFisher Scientific 11791020). Chemically competent TOP10 *E.coli.* (Invitrogen, C404010) were transformed with plasmids containing either Ampicillin or Kanamycin resistance genes and were selected on LB-Agar plates with 100μg/ml Ampicillin or 50μg/ml Kanamycin. *ccdB* resistant chemically competent *E.coli* (Invitrogen, A10460) were transformed with plasmids containing a Gateway cassette (*ccdB*, Chlor.R.) and were selected on LB-Agar plates with 100μg/ml Ampicillin and colonies grown with 100μg/ml Ampicillin and 20μg/ml Chloramphenicol. Plasmid DNA was isolated from bacterial cultures using QIAprep Spin Miniprep Kit (Qiagen 27104) and Sanger sequenced at the DF/HCC DNA Resource Core or GeneWiz. Oligo and dsDNA sequences are listed in Supplemental File 2.

#### *pCFD3-NS* (Addgene #149545)

*pCFD3* (Addgene #49410) (Port *et al.* 2014) was digested with BbsI (Fermentas ER1011) and XbaI (New England Biolabs R0145), which removes the sgRNA scaffold and *Drosophila U6* downstream region, and the backbone was purified using a QIAquick column (28115, Qiagen). A gBlock (IDT) containing two BbsI sites and the *U6* downstream region was inserted into digested *pCFD3* backbone by Gibson assembly.

#### *pCFD5-NS* (Addgene #149546)

*pCFD5* (Addgene #73914) (Port and Bullock 2016) was digested with BbsI (Fermentas ER1011) and XbaI (New England Biolabs R0145), which removes the sgRNA scaffold, *O.s.* Gly tRNA, sgRNA scaffold, and *U6* downstream region. The backbone was purified using a QIAquick column (28115, Qiagen). A gBlock (IDT) containing two BbsI sites and the *U6* downstream region was inserted into the digested *pCFD5* backbone by Gibson assembly. The *D.m.* Gly tRNA sequence remains 5’ to the first BbsI site.

#### *pEntr_PE2* (Addgene #149548)

*PE2* coding sequence was PCR amplified from *pCMV-PE2* (Addgene # 132775). *pEntr* backbone was PCR amplified from *pEntr_D-TOPO* (Invitrogen K240020). *PE2* coding sequence was cloned into *pEntr* backbone by Gibson assembly.

#### *pNos-PE2-attB* (Addgene #149549)

*PE2* coding sequence was PCR amplified from *pCMV-PE2* (Addgene # 132775) and gel purified. *pNos-Cas9-attB* (Ren *et al.* 2013) was digested with XbaI/AvrII (New England Biolabs R0145, R0174) to remove Cas9 sequences and the backbone fragment was gel purified. *PE2* coding sequence was inserted into digested *pNos-attB* by Gibson assembly.

#### *pAct-GW-HygroR* (Addgene #149610)

*Act5c* promoter was amplified from *pAWF* (Murphy lab, unpublished, https://emb.carnegiescience.edu/drosophila-gateway-vector-collection) and gel purified. Backbone was PCR amplified from *pMK33-GW*, using primers that exclude the *Metallothionein* promoter, and gel purified. The *Act5c* fragment was inserted into the *pMK33-GW* backbone by Gibson assembly.

*pUAS-PE2-attB* (Addgene #149550) and *pAct-PE2-HygroR* (Addgene #149552) were generated by Gateway reactions between *pEntr_PE2* and *pWalium10-roe* (Perkins *et al.* 2015) or *pAct-GW-HygroR*, respectively.

To clone the *pCFD3*-PE-*ebony*^*23bpBC*^ expression plasmid, oligos encoding the spacer, scaffold, and extension were inserted into *pCFD3-NS* by ligation. Briefly, *pCFD3-NS* was digested with BbsI and purified on a QIAquick column. Top and bottom oligo pairs encoding either the spacer, scaffold, or extension sequence (Supplemental File 2) were designed such that they had overlapping sticky ends with each other and digested *pCFD3-NS*. Oligo pairs were separately annealed and all were ligated into digested *pCFD3-NS* using T4 ligase (NEB, M0202). See Supplemental File 3 for detailed cloning protocols.

To clone *pCFD3-PE-ebony*^*G111X*^, *pCFD3-PE-white*^*A134X*^, and *pCFD3-PE-forked*^*D111X*^, gBlock (IDT) dsDNA fragments encoding the entire pegRNA were inserted into *pCFD3-NS* by Gibson assembly. Briefly, *pCFD3-NS* was digested with BbsI and purified on a QIAquick column. gBlock fragments were designed such that the pegRNA sequence was flanked by sequence homologous to digested *pCFD3-NS* (Supplemental File 2). For each gene target, a gBlock was inserted into digested *pCFD3-NS* by Gibson assembly. See Supplemental File 3 for detailed cloning protocols.

To clone *pCFD5-PE3-ebony*^*G111X*^, *pCFD5-PE3-white*^*A134X*^, and *pCFD5-PE3-forked*^*D111X*^, two overlapping gBlock (IDT) dsDNA fragments encoding the pegRNA and nicking sgRNA were inserted into *pCFD5-NS* by Gibson assembly. Briefly, *pCFD5-NS* was digested with BbsI and purified on a QIAquick column. gBlock #1 encoded the sgRNA sequence flanked by sequence homologous to *pCFD5-NS* and a partial sequence encoding the *O.s.* Gly tRNA, and gBlock #2 encoded the pegRNA flanked by the *O.s.* Gly tRNA and sequence homologous to *pCFD5-NS* (Supplemental File 2). For each gene target, gBlocks #1&2 were inserted together into digested *pCFD5-NS* by Gibson assembly. See Supplemental File 3 for detailed cloning protocols.

### Cell culture

*Drosophila* S2R+ cells were cultured at 25°C using Schneider’s media (21720-024, ThermoFisher) with 10% FBS (A3912, Sigma) and 50 U/ml penicillin-streptomycin (15070-063, ThermoFisher). S2R+ cells were transfected using Effectene (301427, Qiagen) following the manufacturer’s instructions. S2R+ cells stably expressing PE2 (PT5-PE2) were generated by transfecting *pAct-PE2-HygroR* into the PT5 line (Neumuller *et al.* 2012), which expresses a mCherry-Clic protein trap. PT5 cells were transfected in a 6-well dish at a concentration of 1.8×10^6 cells/ml (2ml total volume). 24 hours after transfection, 200 μg/ml Hygromycin B (Calbiochem 400051-1MU) was added to the media. 5 days after transfection, cells were resuspended and transferred to a T75 flask with fresh media containing 200 μg/ml Hygromycin B. 1 week later, cells were resuspended, centrifuged at 100g for 10min, and resuspended in 3ml fresh media containing 200 μg/ml Hygromycin B. Resuspended cells were transferred serially into each well of a 6-well plate as a dilution series. Visible colonies were resuspended and expanded after ~3 weeks.

Plasmids were transfected into PT5 or PT5-PE2 cells. Briefly, PT5 or PT5-PE2 cells were seeded at 600,000 cells/well of a 24-well plate and transfected with a total of 200 ng plasmid DNA. PT5 cells were transfected with *pAct-Gal4* (unpublished, Dr. Y. Hiromi, National Institute of Genetics, Mishima, Japan), *pUAS-PE2*, pegRNA plasmid, and *pAct-GFP* (aka pLib6.6, unpublished) at a 3:3:3:1 ratio. PT5-PE2 cells were transfected with pegRNA plasmid and *pAct-GFP* at a 3:1 ratio. To increase the chances that GFP+ cells contained prime editing plasmids, we transfected less *pAct-GFP* plasmid relative to the other co-transfected plasmids.

4 days after transfection, GFP+ cells were isolated by fluorescence-activated cell sorting (FACS). Cells were first resuspended in culture media and pipetted into a cell straining FACS tube (352235 Corning) to break up cell clump. 50,000 cells with GFP fluorescence in the 60-80^th^ percentile of fluorescence intensity were sorted on an Aria 561 instrument into a single well of a 96-well plate and incubated at 25°C for 24hr.

5 days after transfection, genomic DNA was isolated from sorted and non-sorted cells using the QuickExtract reagent (Lucigen QE09050). In addition, genomic DNA was isolated from non-transfected PT5 cells as a negative control. Briefly, culture media was removed and replaced with the same volume of QuickExtract reagent. The solution was resuspended by pipetting, transferred to a PCR strip tube, incubated at 65°C for 15min, and then 98°C for 2min.

### Fly culture and crosses

Flies were maintained on standard fly food at 25°C, or at 29°C when noted. Fly stocks were obtained from individual labs or the Bloomington Drosophila Stock Center (BDSC) (indicated with BL#). Stocks used in this study are as follows: *yw* (Perrimon Lab), *yw; Sp hs-hid/CyO* (derived from BL7757), *yw;; TM3,Sb/TM6, Tb* (Perrimon Lab), *ywf* (BL1493), *yv nos-phiC31int; attP40* (BL25709), *yv nos-phiC31int;; attP2* (BL25710), *yw*; *tub-Gal4* (BL5138), *yw*; *Act-Gal4* (BL4414), *yw*; *nos-Gal4* (BL4442), *UAS-emptyVK37* (Bellen lab).

Transgenic flies generated in this study (submitted to the BDSC):

*yw; UAS-PE2,w*+ *attP40* (BL#XXXXX)
*yw;; UAS-PE2,w*+ *attP2* (BL#XXXXX)
*yv; pCFD3-PE-ebony^G111X^,v*+ *attP40* (BL#XXXXX)
*yv; pCFD3-PE-white^A134X^,v*+ *attP40* (BL#XXXXX)
*yv; pCFD3-PE-forked^D111X^,v*+ *attP40* (BL#XXXXX)
*yv; pCFD5-PE3-ebony^G111X^,v*+ *attP40* (BL#XXXXX)
*yv; pCFD5-PE3-white^A134X^,v*+ *attP40* (BL#XXXXX)
*yv; pCFD5-PE3-forked^D111X^,v*+ *attP40* (BL#XXXXX)
*yscv; nos-PE2,v*+ *attP40*
*yv;; nos-PE2,v*+ *attP2*

Fly stocks with multiple transgenes (submitted to the BDSC):

*w*; *Act-Gal4*/CyO; *UAS-PE2,w*+ *attP2* (BL#XXXXX)
*w*; *UAS-PE2,w*+ *attP40*; *Tub-Gal4/TM6b* (BL#XXXXX)
*w*; *nos-Gal4*; *UAS-PE2,w*+ *attP2* (BL#XXXXX)

Transgenic flies were generated by phiC31 integration of attB-containing plasmids into either attP40 or attP2 landing sites. Briefly, plasmid DNA was purified twice on QIAquick columns and eluted in injection buffer (100 μM NaPO4, 5 mM KCl), at a concentration of 200 ng/μl. Plasmid DNA was injected into ~50 fertilized embryos (*yv nos-phiC31int; attP40* or *yv nos-phiC31int;; attP2*) and resulting progeny were outcrossed to screen for transgenic founder progeny. *nos-PE2* and pegRNA insertions were isolated by screening for *vermillion*+ eye color. *UAS-PE2* insertions were isolated by screening for *white*+ eye color.

For PE2 toxicity experiments*, Act-Gal4/CyO* or *tub-Gal4/TM3-Sb* was crossed with either *UAS-empty* (ChrII), *UAS-PE2* (ChrII), or *UAS-PE2* (ChrIII) and progeny were raised at either 25°C or 29°C starting at egg deposition. The frequency of PE2 expressing progeny was determined by counting the number of adult non-balancer progeny and dividing by the total number of flies (# non-balancer/# non-balancer + # balancer).

For somatic editing experiments, *Act>PE2* or *tub>PE2* flies were crossed with pegRNA flies and adult PE2/pegRNA progeny analyzed for mutant phenotypes.

For germ line editing experiments, *nos-PE2* or *nos>PE2* flies were crossed with *pCFD5-PE3-e*^*G111X*^ flies and G1 progeny were crossed with *TM3,e*^*1*^/*TM6b,e*^*1*^. To screen different germ cell PE2 genotypes and temperature conditions, G1 crosses were performed as pools of 10 PE2/pegRNA males or females. G1 crosses were performed as single PE2/pegRNA male or female crosses for optimal conditions (*nos>PE2*, 29°C + h.s.). The phenotypes of G2 progeny were scored as either wild-type or *ebony* (dark cuticle pigment) on a fly dissecting scope. To heat shock G1 larvae, we incubated larvae at 37°C for 1hr in five separate treatments after egg deposition: 24hr, 48hr, 72hr, 96hr, and 120hr.

Focal stack images of adult flies were obtained using a Zeiss Axio Zoom V16 fluorescence microscope and merged using Helicon Focus 7. Images were then processed using Adobe Photoshop CS6.

Fly genomic DNA was isolated by grinding a single fly in 50μl squishing buffer (10 mM Tris-Cl pH 8.2, 1 mM EDTA, 25 mM NaCl) with 200μg/ml Proteinase K (3115879001, Roche), incubating at 37°C for 30 min, and 95°C for 2 minutes. For somatic editing experiments, genomic DNA was collected from adult male flies unless otherwise noted. For germ line editing experiments, genomic DNA was collected from both male and female G2 adult flies.

### Amplicon sequencing

Genomic edit sites were amplified by PCR to yield amplicons for NGS. Briefly, 1μl of S2R+ or fly genomic DNA was used in a PCR reaction using Q5 High-Fidelity DNA Polymerase (NEB M0491L). Primer pairs (Supplemental File 2) were designed to yield amplicons ~200-280 bp in size with the intended editing site located within 100 bp of either the forward or reverse primer. PCR fragments were purified using QIAquick columns (28115, Qiagen) and submitted to the MGH CCIB DNA Core (CRISPR Sequencing), or Genewiz (Amplicon-EZ).

NGS reads were analyzed using CRISPResso2 (version 2.0.38) (Clement *et al.* 2019). To calculate the percent of reads with the precise edit, we used the following parameters: “--prime_editing_pegRNA_spacer_seq”, “--prime_editing_pegRNA_extension_seq”, “--prime_editing_pegRNA_scaffold_sequence”, “--ignore_substitutions”, and “-- discard_indel_reads”. The precise editing frequency was calculated from “CRISPResso_quantification_of_editing_frequency.txt”, for the “Prime-edited” amplicon, as the # unmodified/reads aligned all amplicons. To determine the percent of reads with indels, we ran CRISPResso2 with standard settings and the --ignore_substitutions parameter. The indel frequency was calculated from “CRISPResso_quantification_of_editing_frequency.txt”, as the # modified/# reads_aligned.

For S2R+ and fly experiments involving the edits *ebony*^*G111X*^, *white*^*A134X*^, *forked*^*D111X*^, we specified a quantification window (“-qwc”) that encompasses the region between the pegRNA and nicking sgRNA (spanning the −6 position relative to the pegRNA PAM to the −6 position relative to the sgRNA PAM) (*ebony*: 96-158; *forked*: 97-159; *white*: 112-187).

Fastq files containing amplicon reads will be deposited at the NCBI SRA.

---

**Supplemental Figure 1.**
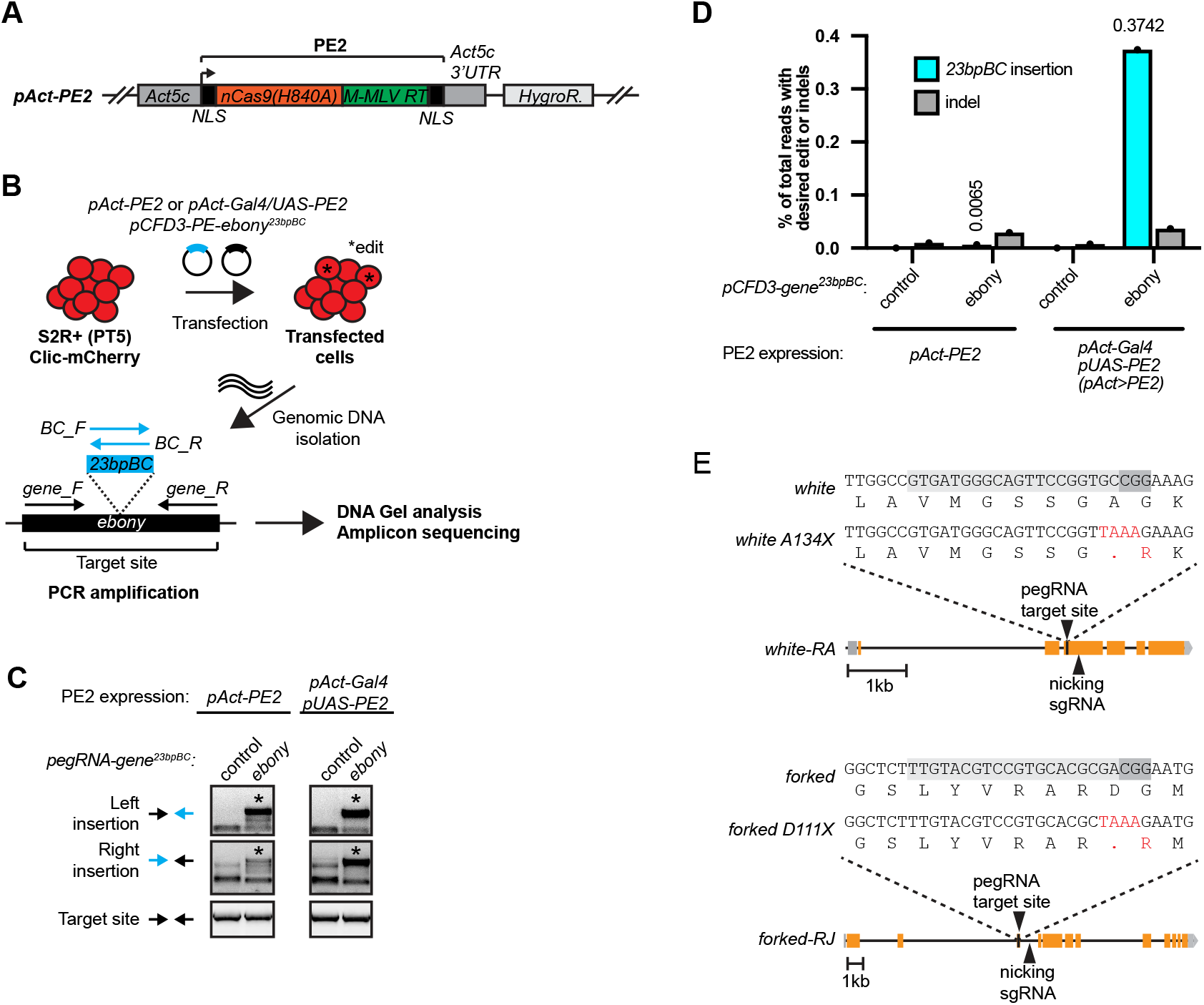
Related to Figure 1. **A.** Diagram of PE2 expression plasmid *pAct-PE2*. **B.** Schematic of experiments to detect and quantify insertion events in transfected S2R+ cells. **C.** DNA gel images of targeted PCR amplification of the insertion site. **D.** Quantification of precise *ebony*^*23bpBC*^ insertion and indel percentage from S2R+ transfection experiments by amplicon sequencing. **E.** *white* and *forked* genomic region showing target site and installed edits (*white*^*A134X*^ and *forked*^*D111X*^).

**Supplemental Figure 2.**
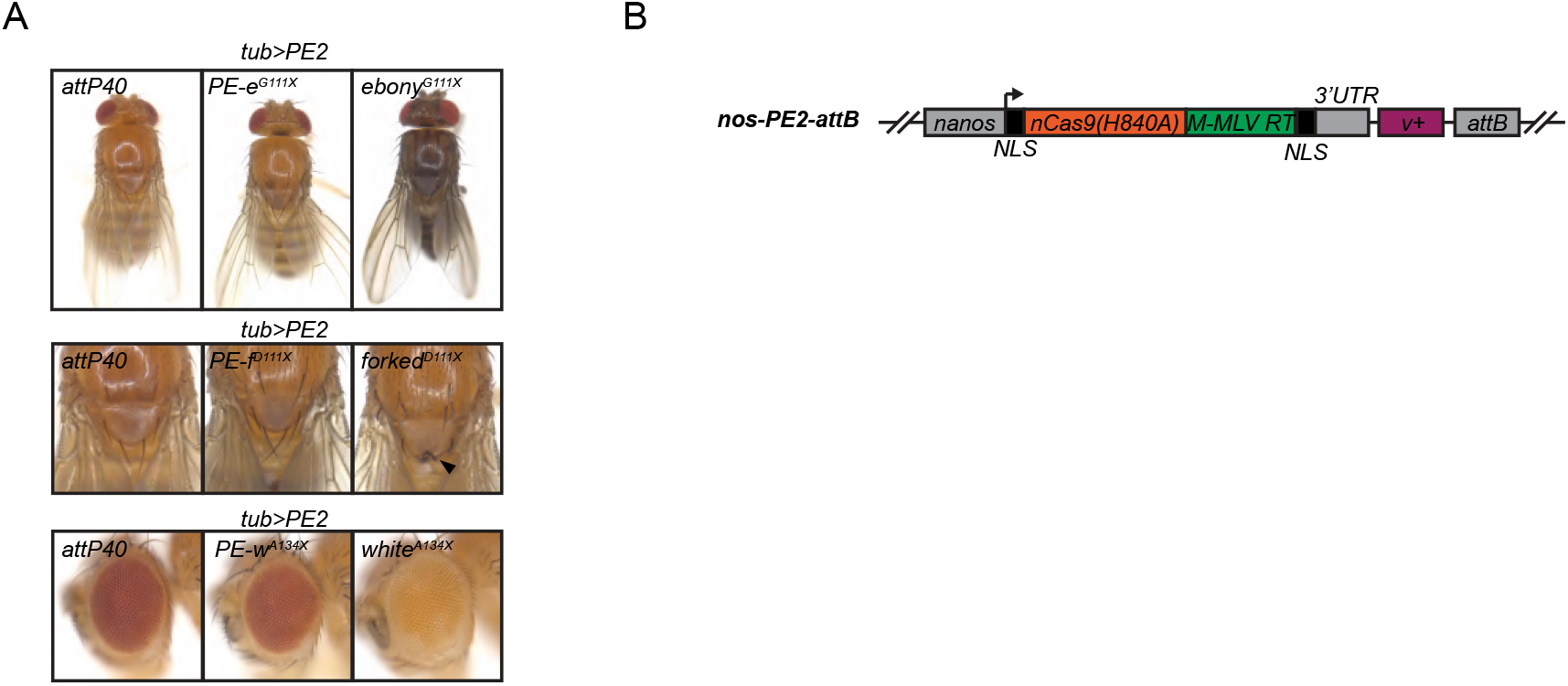
Related to Figure 2. **A.** Images of adult flies with somatic editing using *tub>PE2*. Views of the dorsal side of whole adults (top), scutellum (middle), and eye (bottom). Negative control is *attP40* and classical loss of function allele shown on right. Females shown for editing of *ebony* and *forked*, males shown for *white* editing. **B.** Diagram of PE2 expression transgene *nos-PE2*. *nos, nanos*; NLS, Nuclear localization sequence; 3’ UTR, nanos 3’ UTR; *v+*, *vermillion+* rescue transgene; *attB*, phiC31 recombination site.

**Supplemental File 1.**
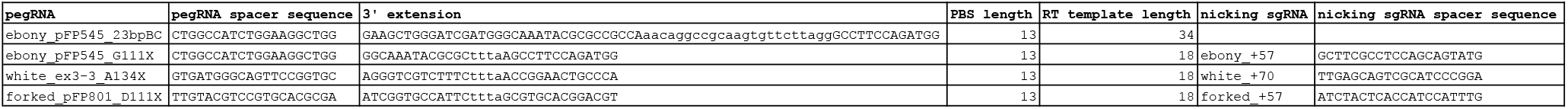
pegRNA and sgRNA sequences.

**Supplemental File 2.**
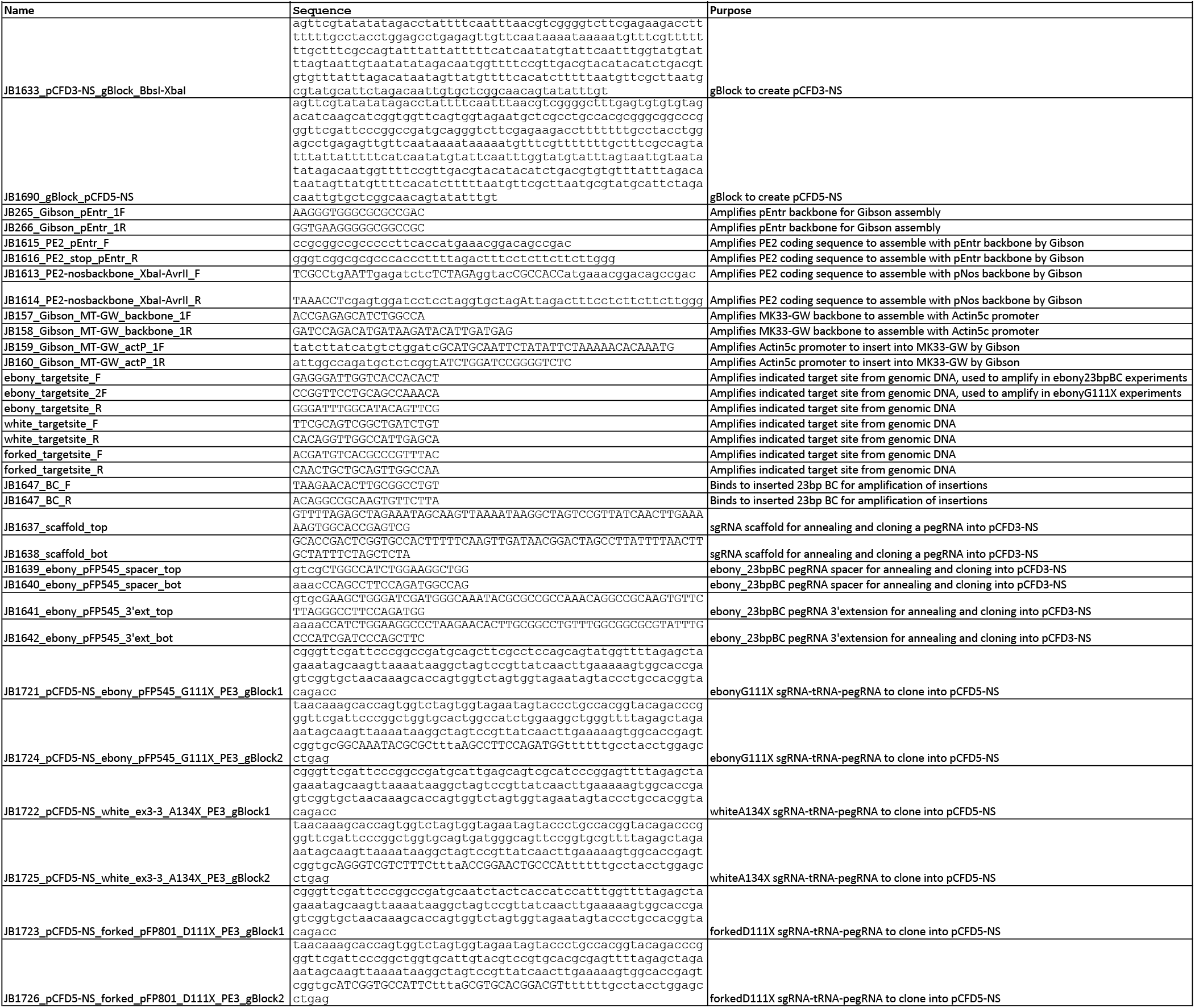
Oligo and dsDNA sequences.

Supplemental File 3. pegRNA design and cloning protocols

Supplemental File 4. Amplicon sequencing data key

## Supplemental File 3

pegRNA cloning for Prime Editing in *Drosophila*, June 2020, Version 1.0

Justin Bosch, Perrimon Lab, Harvard Medical School

### Table of contents

A. Introduction
B. pegRNA and nicking sgRNA design
C. Oligo and dsDNA design
D. Cloning protocol for *pCFD3-NS* using annealed oligos
E. Cloning protocol for *pCFD3-NS* using a dsDNA fragment
F. Cloning protocol for *pCFD5-NS* using two dsDNA fragments

## A. Introduction

These protocols are used to assemble plasmids to express pegRNAs under the control of the *Drosophila U6-3* promoter. pegRNAs are designed to make a precise edit in the genome, and optional nicking sgRNAs are designed to enhance prime editing efficiency (PE3 system). To express pegRNAs and sgRNAs, they are encoded in annealed oligos or dsDNA fragments, and then cloned into one of two empty expression plasmids. *pCFD3-NS* is used for expression of a single pegRNA. *pCFD5-NS* is used for expression of a pegRNA/sgRNA pair. *pCFD3-NS* and *pCFD5-NS* do not contain a sgRNA scaffold (NS = <u>N</u>o <u>S</u>caffold), and are slight modifications of the sgRNA-expression plasmids *pCFD3* and *pCFD5* (Port *et al.* 2014; Port and Bullock 2016). *pCFD3-NS* and *pCFD5-NS* contain an *attB* site for phiC31 integration and a *vermillion+* marker to select transgenic flies.

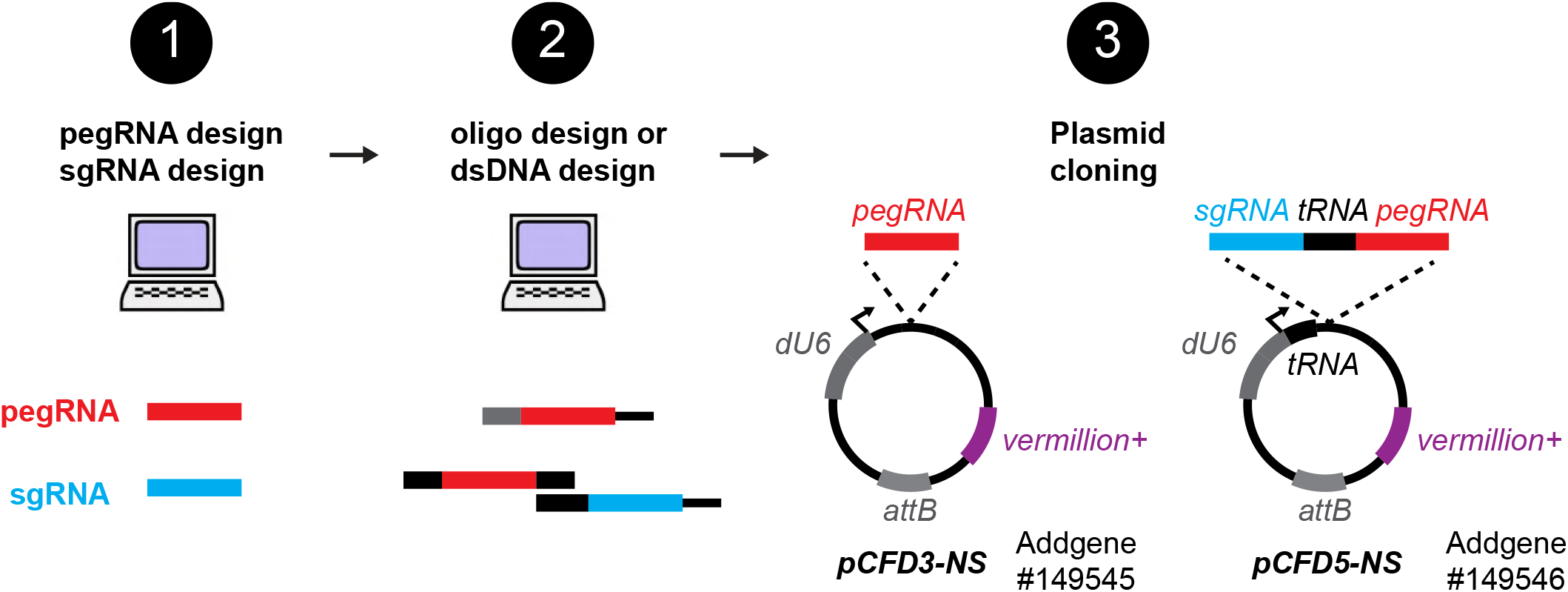

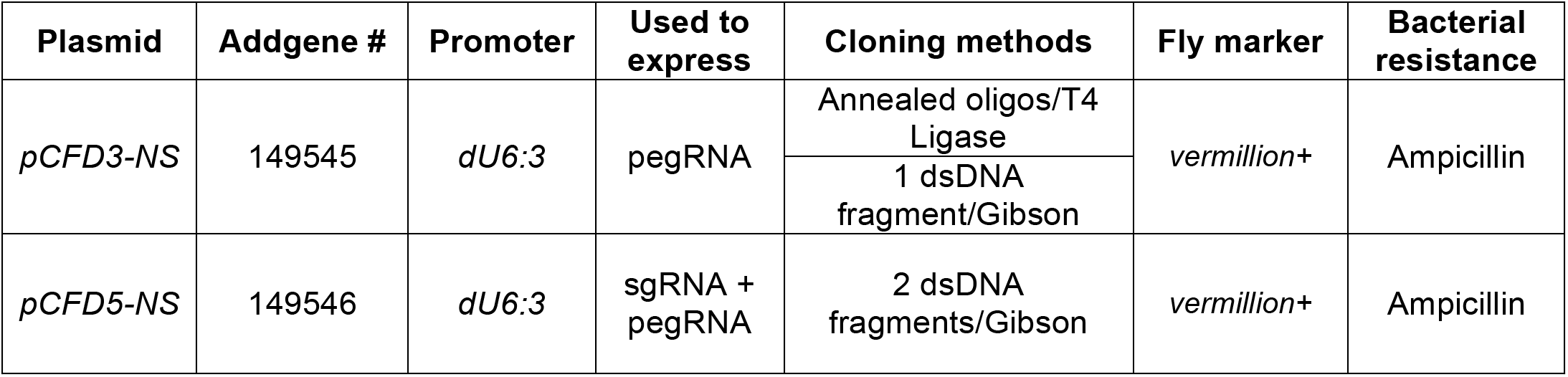
Summary of pegRNA-expression plasmids.

## B. pegRNA and nicking sgRNA design

Automatic design (recommended):

PrimeDesign (Hsu *et al.* 2020): http://primedesign.pinellolab.org/
pegFinder (Chow *et al.* 2020): http://pegfinder.sidichenlab.org/

Manual design (optional):

1. Create wild-type (WT) and edited sequence files for annotation
2. WT sequence - select a pegRNA spacer near the desired edit, ensuring the edit is 3’ to nick site.
3. Edited sequence - annotate the primer binding site (PBS) by selecting ~13bp 5’ to the nick site.
4. Edited sequence - annotate the reverse transcribed (RT) region by selecting ~13-18bp 3’ to nick site.
5. Edited sequence – The reverse complement of the PBS-edit-RT sequence is the pegRNA 3’ extension.
6. WT sequence - select a sgRNA target on the non-edited strand between +40 and +90 from the pegRNA nick.

Notes:

- Avoid starting pegRNA 3’ extension with a “C”.
- Edits or silent mutations that affect the PAM or pegRNA spacer sequence increase efficiency.
- Use a shorter RT sequence if region has high G:C content.

Example pegRNA and nicking sgRNA design:

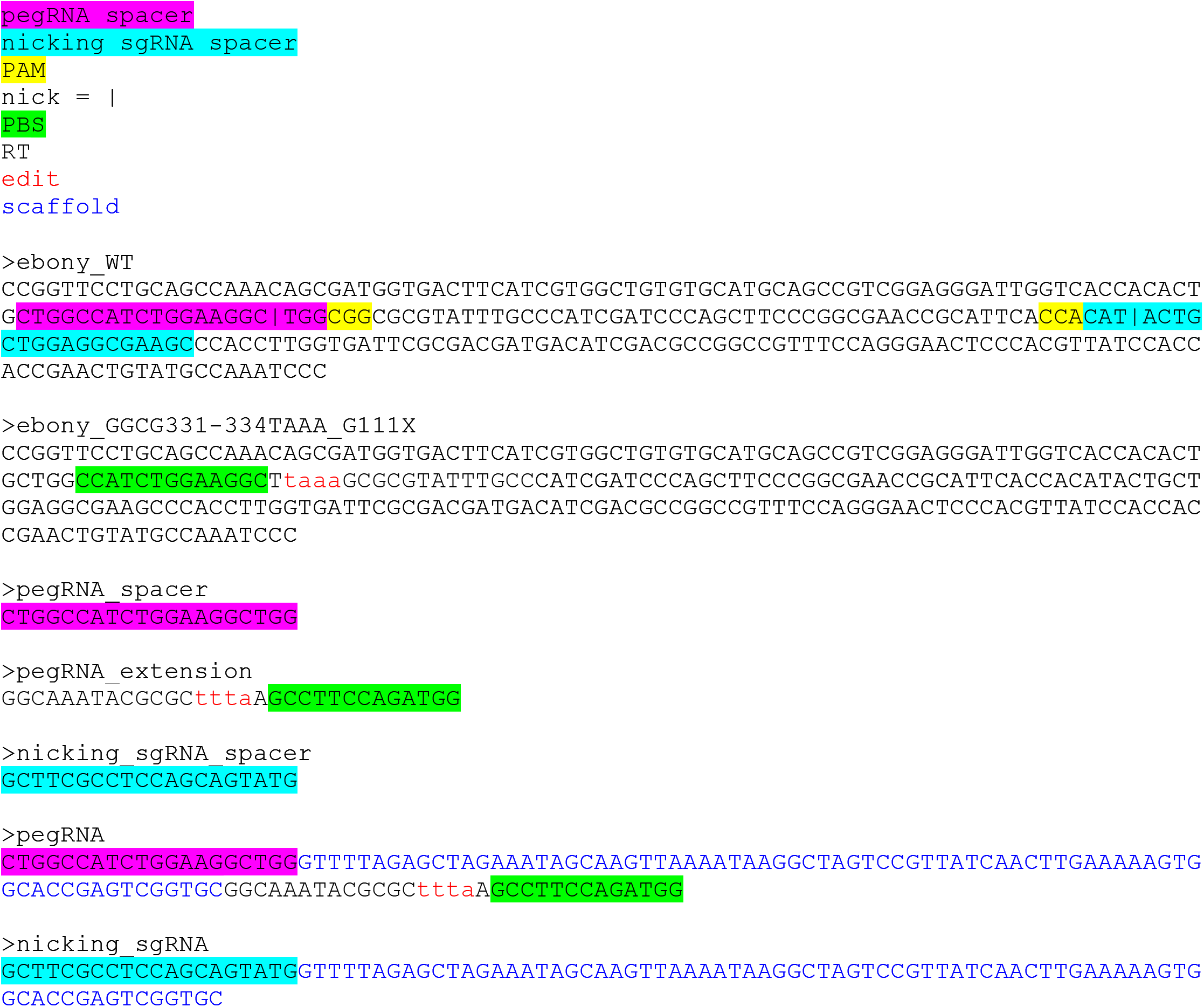

## C. Oligo and dsDNA design

### C1. For cloning into pCFD3-NS by T4 ligation (single pegRNA) (See section D)

Order oligos with overhangs (5’ lowercase sequence)

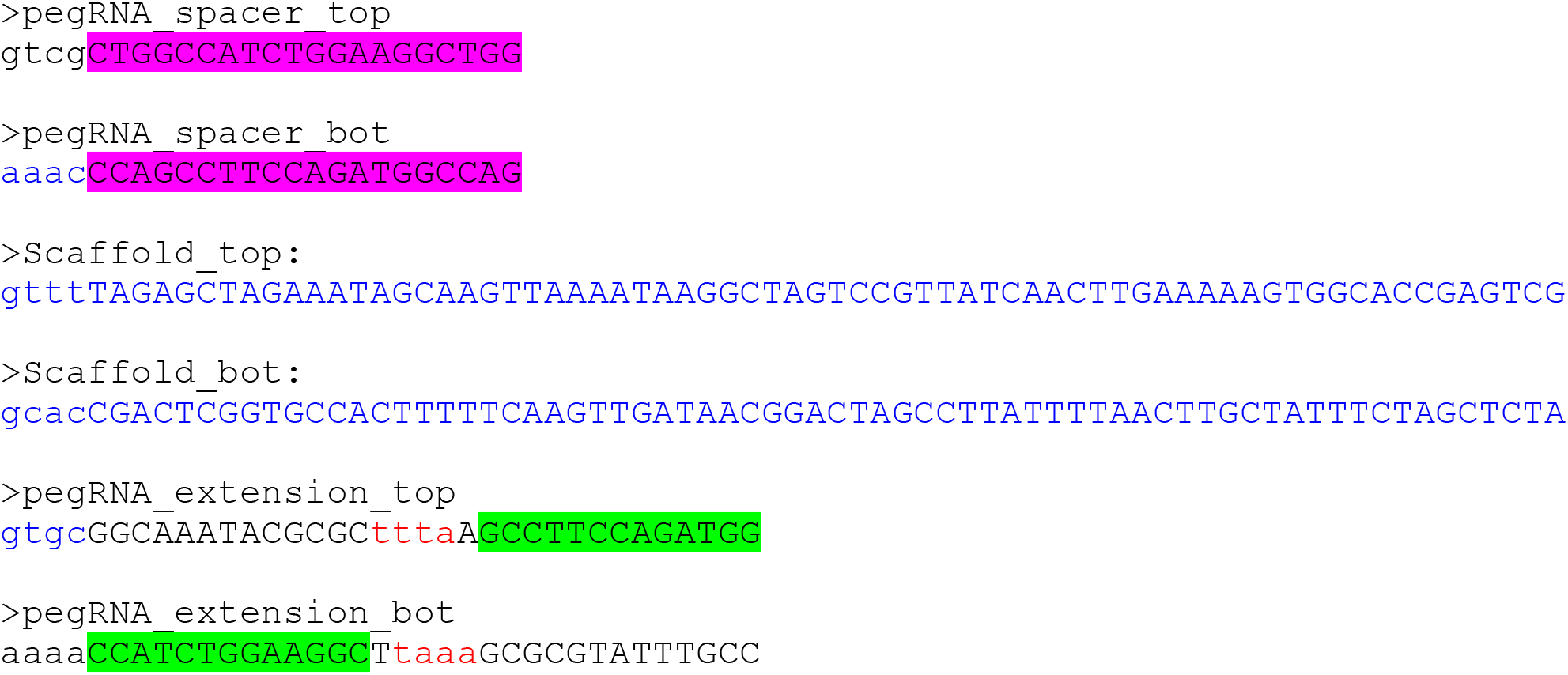

Annealed oligos:

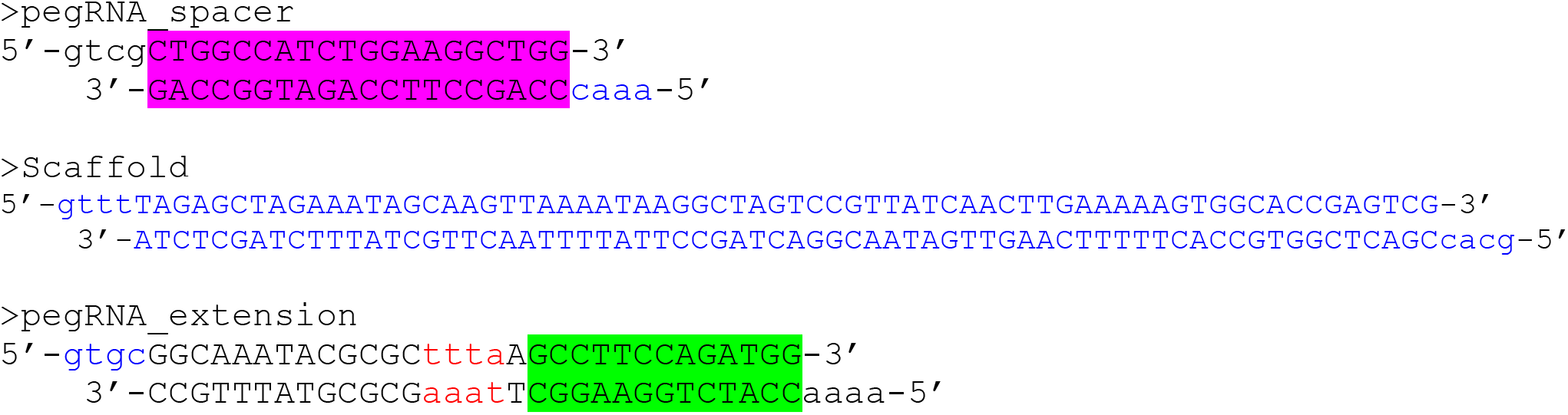

Cloning:

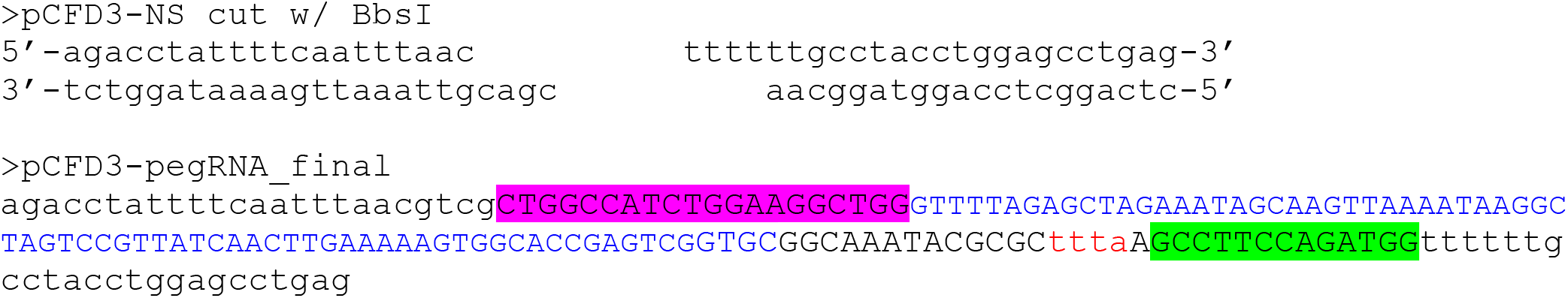

### C2. dsDNA to clone into pCFD3-NS by Gibson assembly (single pegRNA) (See section E)

Append homology arms (black, lowercase) to pegRNA that overlap with pCFD3-NS cut w/ BbsI.

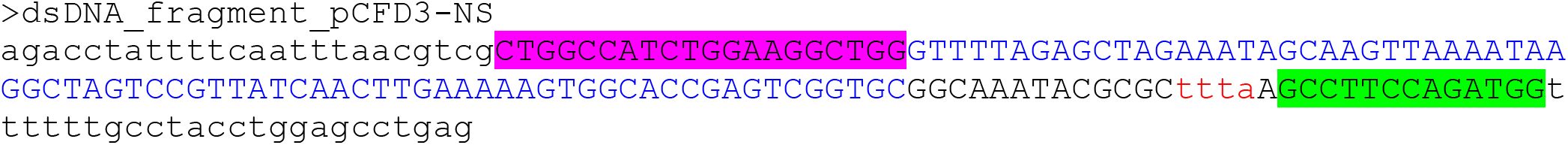

Cloning:

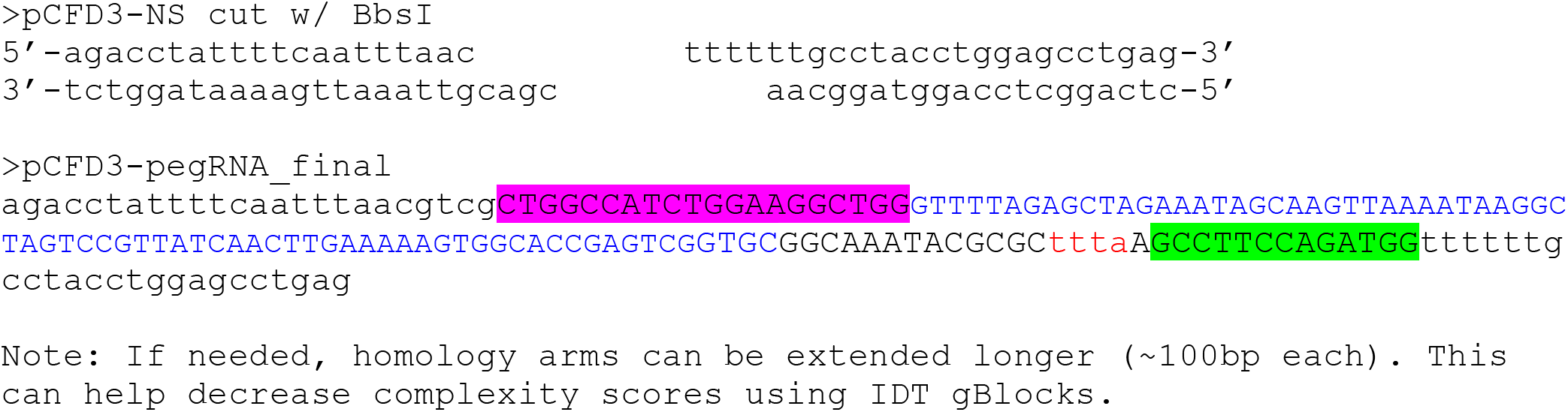

### C3. dsDNAs to clone into pCFD5-NS by Gibson assembly (nicking sgRNA and pegRNA) (See section F)

Append homology arms (black, lowercase) to nicking sgRNA and pegRNA that overlap with pCFD3-NS cut w/ BbsI and encode rice Os-tRNA^Gly^ (lowercase, italic)

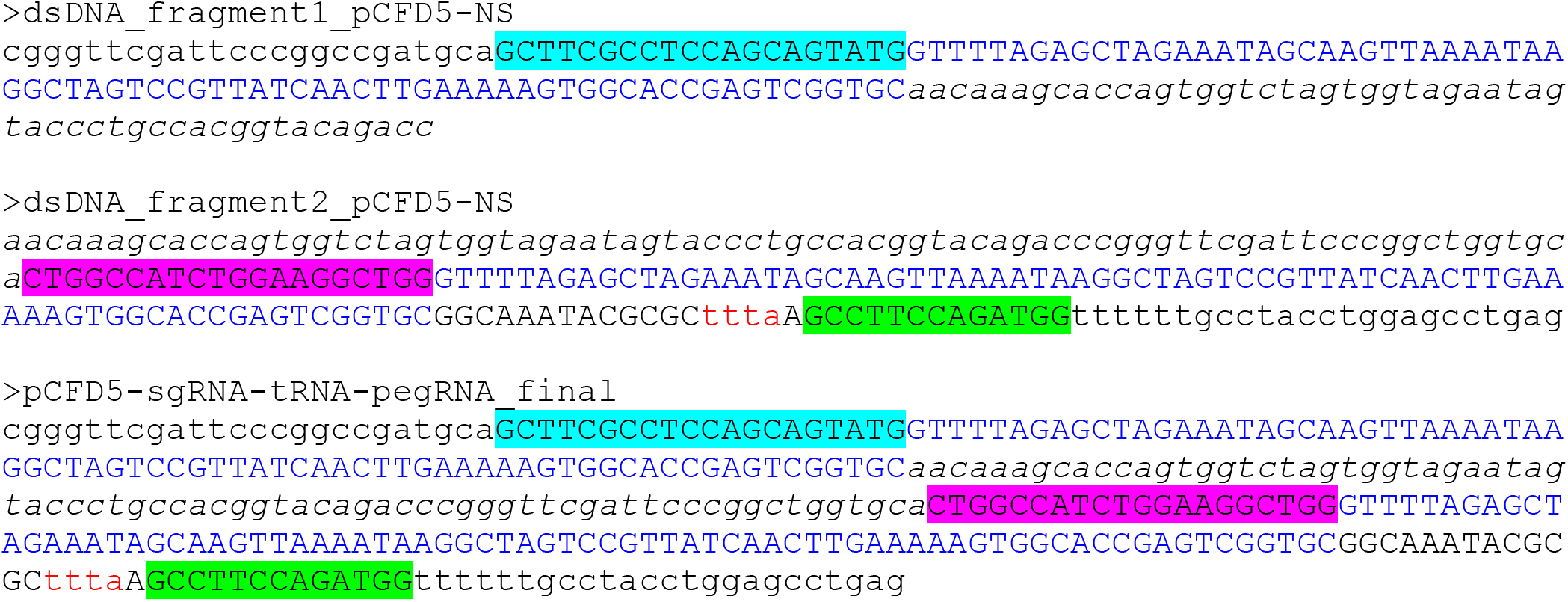

## D. Cloning protocol for *pCFD3-NS* (Addgene # 149545) using annealed oligos

### D1. Design pegRNA and order oligos (see Sections B&C)

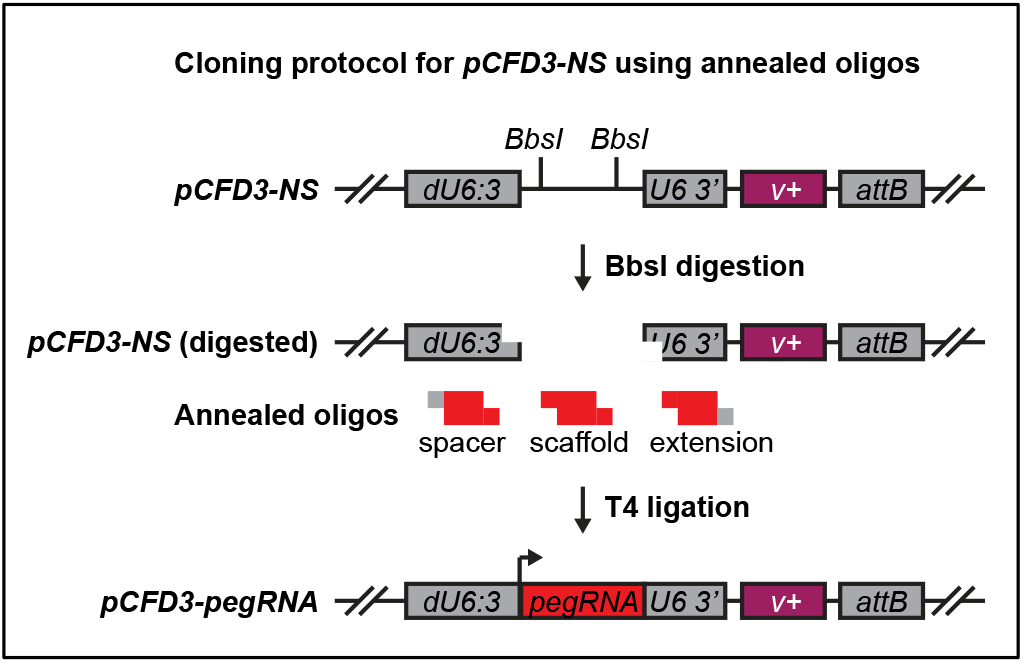

### D2. Digest/dephosphorylate *pCFD3-NS*

5μg pCFD3-NS

3μl BpiI (cuts BbsI) (Fermentas, FD1014)

3μl FastAP (Fermentas, EF0651)

6μl 10x FastDigest Buffer

<u>Xμl H20</u>

60ul total

### D3. Gel-purify digested *pCFD3-NS* backbone (~6.2kb)

### D4. Phosphorylate and anneal each pair of oligos in PCR tubes

1μl Top oligo (100μM)

1μl Bottom oligo (100μM)

1μl 10x T4 Ligation buffer (NEB, B0202S)

6.5μl H20

<u>.5μl T4 PNK (NEB, M0201)</u>

10μl total

37°C for 30min, 95°C for 5min, then ramp down to 25°C at 5°C/min

### D5. Dilute annealed/phosphorylated oligos 1:200 in H20

### D6. Ligate annealed oligos into digested *pCFD3-NS*

Xμl digested pCFD3-NS (50ng)

1μl **spacer** diluted annealed oligo

1μl **scaffold** diluted annealed oligo

1μl **3’ extension** diluted annealed oligo

1.5μl 10x T4 Ligation Buffer (NEB, B0202S)

Xμl H20

<u>1μl T4 DNA ligase (NEB, M0202)</u>

15μl total

Incubate reaction at room temperature for 30min.

### D7. Transform ligation into competent cells and grow colonies on LB-agar Ampicillin plates

### D8. (Optional) Colony PCR to identify candidate pegRNA plasmids

pCFD3genoF ACGTTTTATAACTTATGCCCCTAAG

pCFD3genoR GCCGAGCACAATTGTCTAGAATGC

Uncut backbone = 490bp

Correct insert = 638bp (depends on pegRNA length)

### D9. Culture colonies with LB + Ampicillin and sequence confirm plasmids

pCFD3seqF ACCTACTCAGCCAAGAGGC

## E. Cloning protocol for *pCFD3-NS* (Addgene # 149545) using a dsDNA fragment

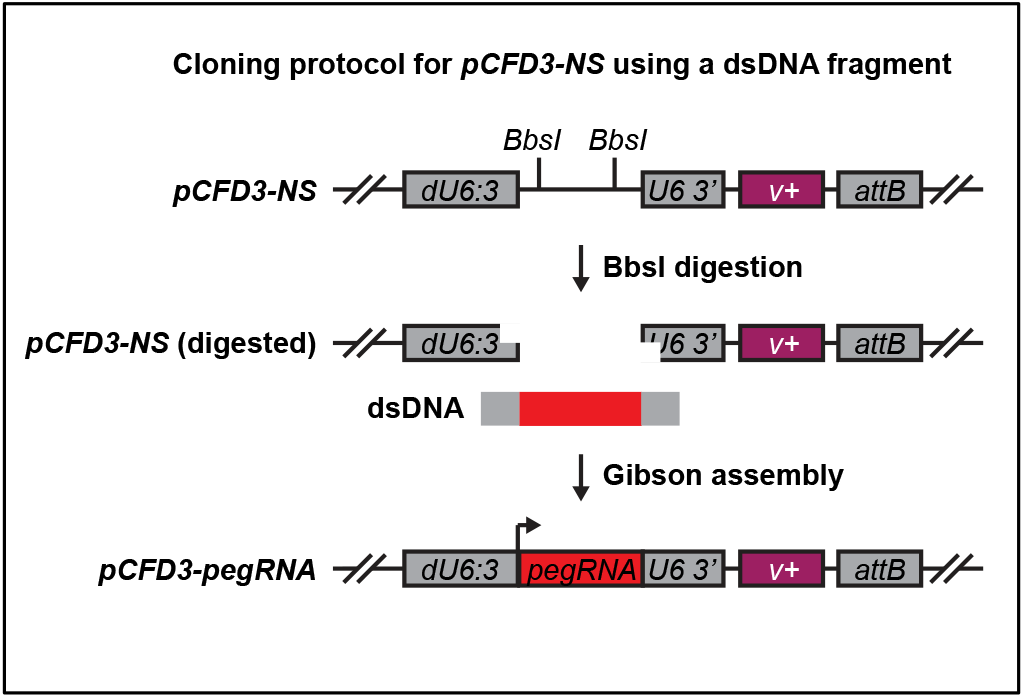

### E1. Design pegRNA and order dsDNA fragment (see Sections B&C)

### E2. Digest/dephosphorylate plasmid

5μg pCFD3-NS

3μl BpiI (cuts BbsI) (Fermentas, FD1014)

3μl FastAP (Fermentas, EF0651)

6μl 10x FastDigest Buffer

<u>Xμl H20</u>

60ul total

### E3. Gel-purify digested *pCFD3-NS* backbone (~6.2kb)

### E4. Gibson assembly

Xμl digested pCFD3-NS (50ng)

Xμl dsDNA fragment (5ng)

2.5μl Gibson master mix (NEB, E2611)

<u>Xμl H20</u>

5μl total

Incubate reaction at 50°C for 30min.

### E5. Transform ligation into competent cells and grow colonies on LB-agar Ampicillin plates

### E6. (Optional) Colony PCR to identify candidate pegRNA plasmids

pCFD3genoF ACGTTTTATAACTTATGCCCCTAAG

pCFD3genoR GCCGAGCACAATTGTCTAGAATGC

Uncut backbone = 490bp

Correct insert = 638bp (depends on pegRNA length)

### E7. Culture colonies with LB + Ampicillin and sequence confirm plasmids

pCFD3seqF ACCTACTCAGCCAAGAGGC

## F. Cloning protocol for *pCFD5-NS* (Addgene # 149546) using two dsDNA fragments

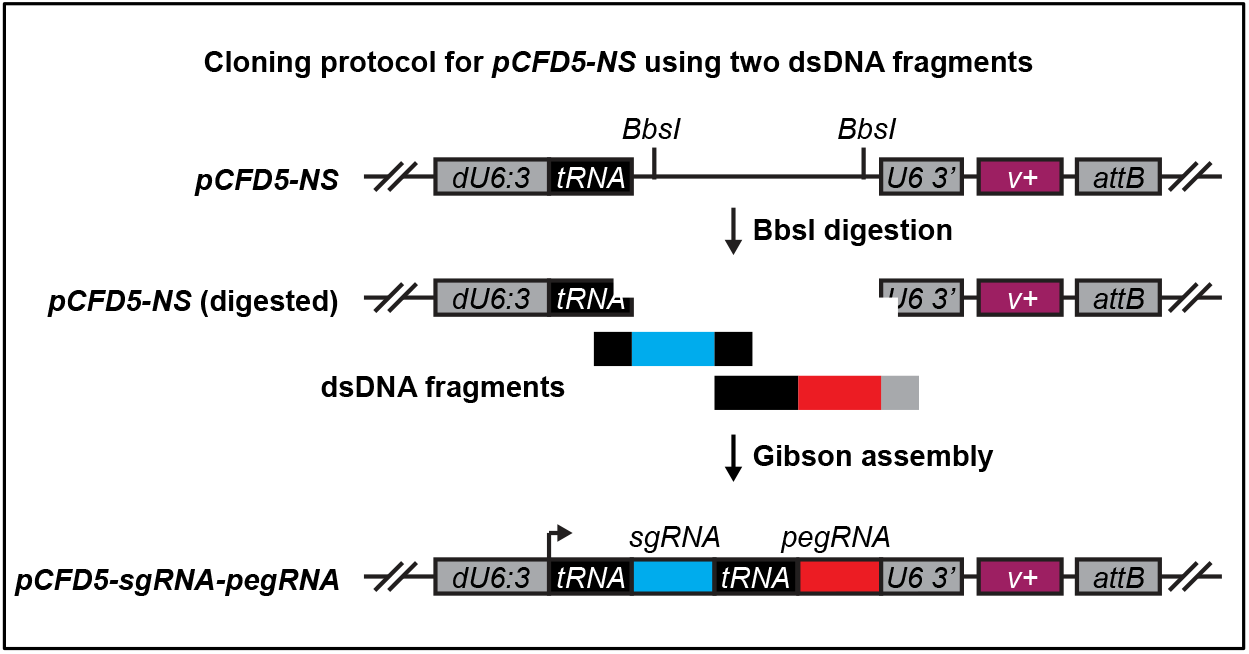

### F1. Design pegRNA and nicking sgRNA, and order dsDNA fragments (see Sections B&C)

### F2. Digest/dephosphorylate plasmid

5μg *pCFD5-NS*

3μl BpiI (cuts BbsI) (Fermentas, FD1014)

3μl FastAP (Fermentas, EF0651)

6μl 10x FastDigest Buffer

<u>Xμl H20</u>

60ul total

### F3. Gel-purify digested *pCFD5-NS* backbone (~6.3kb)

### F4. Gibson assembly

Xμl digested pCFD5-NS (50ng)

Xul dsDNA fragment 1 (5ng)

Xul dsDNA fragment 2 (5ng)

2.5μl Gibson master mix (NEB, E2611)

<u>Xμl H20</u>

5ul total

Incubate reaction at 50°C for 30min.

### F5. Transform ligation into competent cells and grow colonies on LB-agar Ampicillin plates

### F6. (Optional) Colony PCR to identify candidate pegRNA plasmids

pCFD3genoF ACGTTTTATAACTTATGCCCCTAAG

pCFD3genoR GCCGAGCACAATTGTCTAGAATGC

Uncut backbone = 587bp

Correct insert = ~846bp (depends on pegRNA length)

### F7. Culture colonies with LB + Ampicillin and sequence confirm plasmids

pCFD3seqF ACCTACTCAGCCAAGAGGC

